# Entropy-driven translocation of disordered proteins through the Gram-positive bacterial cell wall

**DOI:** 10.1101/2020.11.24.396366

**Authors:** David K. Halladin, Fabian E. Ortega, Katharine M. Ng, Matthew J. Footer, Nenad S. Mitić, Saša N. Malkov, Ajay Gopinathan, Kerwyn Casey Huang, Julie A. Theriot

**Affiliations:** Department of Microbiology & Immunology, Stanford University School of Medicine, Stanford, CA 94305; Department of Biochemistry and Howard Hughes Medical Institute, Stanford University School of Medicine, Stanford, CA 94305; Department of Bioengineering, Stanford University, Stanford, CA 94305; Department of Biology and Howard Hughes Medical Institute, University of Washington, Seattle, WA 98185; Faculty of Mathematics, University of Belgrade, Belgrade, Serbia; Department of Physics, University of California, Merced, CA 95343; Chan Zuckerberg Biohub, San Francisco, CA 94158

## Abstract

Cells across all kingdoms of life actively partition molecules between discrete cellular compartments. In Gram-positive bacteria, a thick and highly cross-linked peptidoglycan cell wall separates the bacterial membrane from the extracellular space, imposing a barrier that must be crossed by proteins whose functions require that they be exposed on the bacterial cell surface^1,2^. Some surface-exposed proteins, such as the *Listeria monocytogenes* actin nucleation-promoting factor ActA^3^, remain associated with the bacterial membrane yet somehow thread through tens of nanometers of dense, cross-linked cell wall to expose their N-terminus on the outer surface^4,5^. Here, we show that entropy can drive the translocation of disordered transmembrane proteins through the Gram-positive cell wall. We develop a physical model predicting that the entropic constraint imposed by a thin periplasm is sufficient to drive translocation of an intrinsically disordered protein like ActA across a porous barrier similar to the cell wall. Consistent with this scenario, we demonstrate experimentally that translocation depends on both the dimensions of the cell envelope and the length of the disordered protein, and that translocation is reversible. We also show that disordered regions from eukaryotic nuclear pore complex proteins are capable of entropy-driven translocation through Gram-positive cell walls. These observations suggest that entropic forces alone, rather than chaperones or chemical energy, are sufficient to drive translocation of certain Gram-positive surface proteins for exposure on the outer surface of the cell wall.

---

Surface-exposed proteins are used by both commensal and pathogenic bacteria to mediate a range of processes that are essential for survival within a mammalian host^6,7^. These host-microbe interactions often involve binding of bacterial proteins to host factors that are too large to diffuse through the nanometer-scale pores of the bacterial cell wall^8^. Therefore, such bacterial surface proteins must navigate tens of nanometers through the peptidoglycan to reach the outer cell surface in order to interact with their host-encoded partners^9,10^. A particular topological challenge is faced by membrane-anchored Gram-positive surface proteins. In their mature form, these proteins must span all the cellular compartments starting from the cytoplasm, reaching across the membrane, traversing the periplasmic-like space subjacent to the wall^11^, and then threading through the thick peptidoglycan to expose their functional domains on the far side of the wall.

One well-characterized protein in this category is the virulence factor ActA from *Listeria monocytogenes^3^*. This food-borne pathogen uses a form of bacterial-directed actin-based motility to propel itself through the cytoplasm of a host cell and to spread from cell to cell in an infected mammalian host^12^. ActA is the only bacterial factor required for host cell actin assembly^13,14^, which it accomplishes through direct interaction with the host-encoded actin nucleation factor Arp2/3 complex and host-encoded actin elongation factor VASP in the host cell cytoplasm^15^. ActA is a large (~600 amino acid) protein that is intrinsically disordered^16^. Its experimentally determined Stokes radius of 8 nm is substantially larger than the size of the pores in the Gram-positive cell wall, which limit the passive transport of macromolecules to a cutoff radius of ~2-3 nm^8^. However, it has been shown that intrinsically disordered proteins may tunnel through very small pores, probably by reptation^17^. It is therefore plausible that ActA can similarly thread through the cell wall pores.

Some form of free energy is generally required for vectorial transport of macromolecules across the barriers that separate cellular compartments, often ATP hydrolysis or transmembrane potential^18^. Once a protein has exited the cytoplasmic space, there is no obvious source of chemical free energy to drive directional translocation across the cell wall. How then does ActA translocate its N-terminal domain while still remaining anchored to the cell membrane? Truncation of ActA before the C-terminal transmembrane domain results in secretion of the protein into the extracellular medium^5^, suggesting that there are no strong binding associations between ActA and the cell wall that could either help or hinder its translocation. In Gram-negative bacteria, chaperones shuttle proteins and other cargo from the inner membrane to the outer membrane^19^. However, no such chaperone-mediated transport mechanism has been described for protein export through the Gram-positive cell wall.

Here, we explore the possibility that confinement of a disordered protein with a large hydrodynamic radius, such as ActA, within the periplasmic-like space results in an entropic force that drives protein translocation through the cell wall in the absence of any specific chaperones or other biological machinery. Most ActA molecules were extractable from the cell wall in LB and Welshimer's Broth (WB) media (Fig. 1a), and only ActA was extracted from wild-type cells by SDS boiling, indicating its unique translocated topology (Fig. 1b). To determine whether entropic forces are sufficient to drive ActA translocation, we developed a physical model for the confinement of a disordered polymer in the periplasm and cell wall (Supplementary Text). The model uses three parameters to characterize the dimensions of the bacterial cell envelope: cell wall thickness, *w;* periplasmic thickness, *P;* and wall pore radius, *R*. We treat the polymer as a self-avoiding chain with a persistence length estimated from the experimentally determined Stokes radius of ActA^16^. Our model predicts that, due to the free energy cost of crossing the cell wall, short polymers will remain in the periplasm even for polymer lengths at which they experience some confinement, because confinement of a part of the polypeptide chain within a narrow cell wall pore adds significantly to the entropic penalty. However, for any given *w*, *P*, and *R*, there is a critical length beyond which the entropic cost of periplasmic confinement can be relieved by extending through the cell wall and accessing the unconfined exterior (Fig. 1c,d, Supplementary Text). The length of ActA is greater than the predicted critical length for a wide range of plausible values of *w, P*, and *R* (Extended Data Fig. 1). Thus, in theory, entropy alone should be sufficient to drive the translocation of an ActA-like polymer through a porous barrier with dimensions that resemble the Gram-positive cell wall, resulting in a final configuration in which the C-terminus of the protein remains anchored to the bacterial membrane while the large N-terminal domain is available to interact with host cell binding partners.

**Figure 1:**
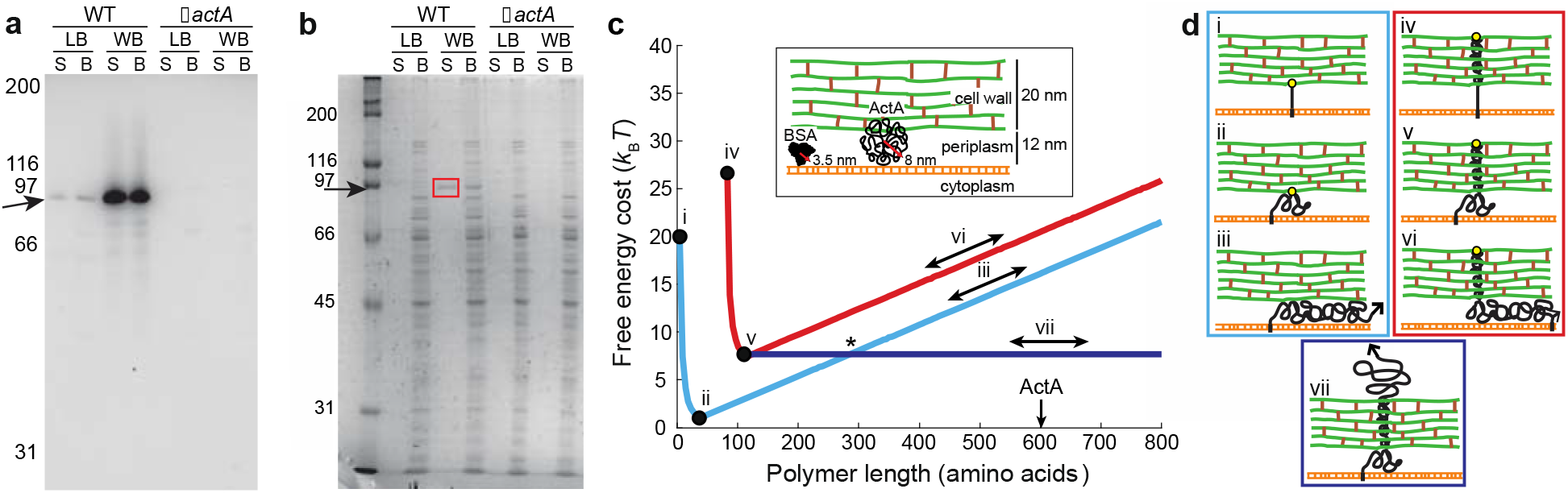
An entropy-based model predicts the navigation of a disordered protein across a Gram-positive cell wall. a) Western blotting of SDS-treated (S) and mechanically disrupted cells through bead-beating (B) demonstrated expression of ActA (arrow) in both LB and Welshimer's Broth (WB) for wild-type (WT) but not in an Δ*actA* mutant. In both growth media, comparable amounts of ActA were recovered from SDS and mechanically disrupted cells, indicating that essentially all ActA is extractable from the cell wall. b) Coomassie staining revealed that cytoplasmic proteins are released after mechanical disruption, whereas only ActA (arrow) was extracted by SDS boiling in wild-type cells (red box), indicating its unique translocated topology. c) Free energy cost for a confined self-avoiding polymer, relative to a polymer in solution, plotted as a function of polymer length using a cell wall thickness of 20 nm, periplasmic thickness of 12 nm, and wall pore radius of 3 nm (see also Supplementary Text and Extended Data Fig. 1). Blue curve: polymer confined to the periplasm; red curve: polymer in which 78 amino acids extend through a pore into the cell wall and all additional amino acids are in the periplasm; horizontal blue line: polymer in an extended state in which 115 amino acids are used to cross the periplasm and cell wall, and all additional amino acids are beyond the outer edge of the cell wall. The asterisk marks the critical length beyond which it is energetically favorable for the polymer to extend through the cell wall. Insert: schematic of the cell surface that includes the relative sizes of ActA and bovine serum albumin (BSA, PDB ID 1E7I), which have similar molecular weights but very different Stokes radii. d) Schematic depicting polymer topology in different states. States i, ii, and iii (blue box) correspond to the blue curve in (**c**), states iv, v, and vi (red box) correspond to the red curve in (**c**), and state vii (fully translocated) corresponds to the navy curve in (**c**). Yellow circles signify constraints where one point in the polymer is fixed to either the inner surface of the cell wall (states i and ii) or the outer surface (states iv and v). Arrows denote topologies of variable polymer length (states iii, vi, and vii) and correspond to the portions of the curves in (**c**) labeled with double-headed arrows.

Our model implicitly assumes that the transition between an extended and periplasmic state is reversible; in this case, polymer topology would be expected to re-equilibrate if the polymer length were to change after translocation (Fig. 2a,b). For a range of parameters corresponding to critical lengths ranging from ~200 to 600 amino acids (Extended Data Fig. 1), our entropic model predicts a relatively sharp sigmoidal transition between a state in which the majority fraction of ActA molecules are fully extended to a state in which the majority fraction of molecules are retained within the periplasm (Fig. 2a). To determine whether truncating full-length ActA below a critical length does indeed result in the re-equilibration of ActA to a periplasmic state, we engineered a series of ActA constructs with an internal TEV protease recognition site positioned at various locations (Fig. 2c). To detect ActA translocation, we used an antibody raised against the VASP-binding domain of ActA, which is normally exposed on the external surface of the bacterial cell wall to bind host proteins^15^. All TEV-site constructs were designed with the protease cleavage site N-terminal of this epitope, so that the epitope would remain intact and anchored to the bacterial membrane following protease cleavage. All constructs displayed comparable cleavage efficiency after treatment with TEV protease, and the cleaved ActA molecules showed no apparent differences in stability (Fig. 2d). Because antibodies are too large to diffuse through the pores in the cell wall^8^, we used immunofluorescence microscopy to differentiate between ActA molecules extended through the cell wall and those confined in the periplasm. We observed that the removal of 166 or more amino acids from ActA resulted in a significant decrease in antibody labeling, whereas shorter truncations had little to no effect (Fig. 2e and Extended Data Fig. 2). We also constructed genetic truncations by removing 100 or 200 amino acids from the N-terminus of ActA (Methods, Extended Data Fig. 3). Consistent with our TEV data, the 100-amino acid truncation exhibited similar antibody labeling to full-length ActA while labeling of the 200-amino acid truncation decreased significantly (Fig. 2f), to a comparable degree as our TEV-166 construct. Thus, truncating ActA past a critical length results in a decrease in the fraction of molecules extending through the cell wall into the extracellular milieu, and ActA passage through the cell wall is reversible.

**Figure 2:**
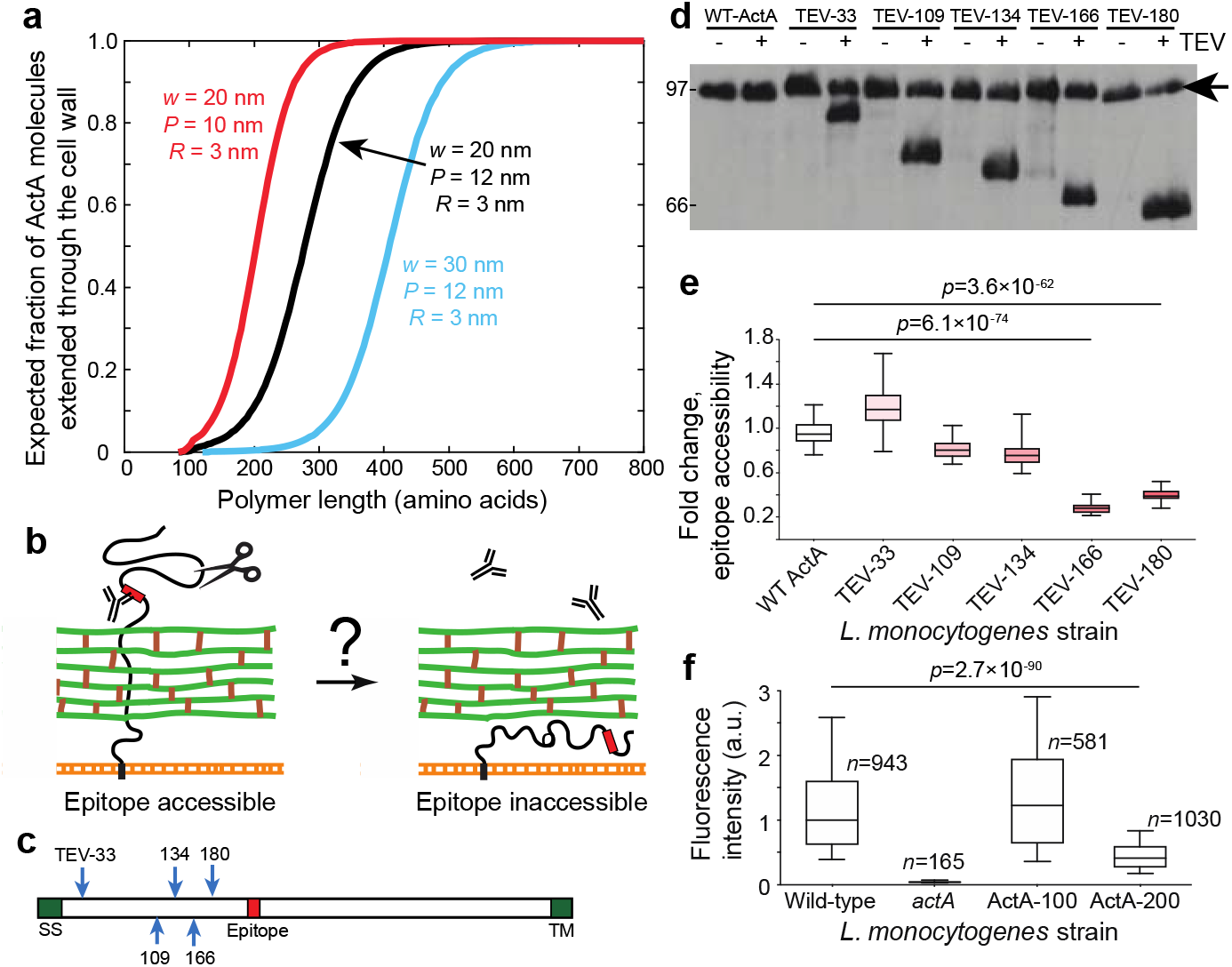
Truncating ActA past a critical length results in a net relocation of ActA molecules into the periplasm. a) Expected fraction of ActA molecules in an extended state as a function of polymer length (Supplementary Text) for various values of cell wall thickness *w*, periplasmic thickness *P*, and wall pore radius *R*. b) Following cleavage by the TEV protease, an ActA molecule can remain extended or transition to a periplasmic state. c) Schematic of ActA showing the position of the signal sequence (SS), transmembrane domain (TM), TEV cleavage sites (arrows), and antibody recognition site. d) Western blotting using an antibody against the proline-rich repeat region of ActA, for strains carrying each construct before (-) and after (+) treatment with TEV protease. Arrow indicates full length ActA. e) The fold change in immunofluorescence intensity corresponding to epitope accessibility comparing untreated and TEV protease-treated populations was significant only for cleavage of 166 or 180 amino acids. Fluorescence intensities were normalized so that the median of the wild-type strain was 1. The box shows 25^th^ and 75^th^ percentiles with the median as a horizontal line, and whiskers are 10^th^ and 90^th^ percentiles. The shades of red indicate the truncation length. For untreated, *n*=515, 541, 382, 508, 277, 505 cells; for TEV-treated, *n*=384, 472, 452, 488, 667, 625 cells. f) Truncation of 200, but not 100, amino acids from the C-terminus of ActA significantly reduced immunofluorescence intensity. Essentially no labeling was detected in an *actA* mutant. Fluorescence intensities were normalized so that the median of the wild-type strain was 1. The box shows 25^th^ and 75^th^ percentiles with the median as a horizontal line, and whiskers are 10^th^ and 90^th^ percentiles. Blue shading indicates confidence intervals for the median.

To determine whether differences in cell surface architecture affect the length dependence of polymer translocation, we substantially decreased cell wall thickness by deleting the *walI* gene (Fig. 3a,b), which encodes a regulator of the WalRK two-component system that coordinates wall synthesis with growth and division^20,21^. We observed similar antibody labeling upon expression of full-length ActA or the 100- or 200-amino acid genetic truncations (Fig. 3c), indicating that unlike in wild-type cells (Fig. 2f) the 200-amino acid truncation was able to translocate. We also found that the cell wall became thicker during growth in WB medium (Fig. 3a,b); the thicker wall would impose a larger entropic barrier, increasing the critical length for entropic translocation. Now, labeling of full length ActA was significantly higher than both the 100- and 200-amino acid truncations (Fig. 3d), indicating that the 100-amino acid truncation was compromised in its ability to translocate across the thicker wall, consistent with the hypothesis that the critical length had increased as predicted by our model.

**Figure 3:**
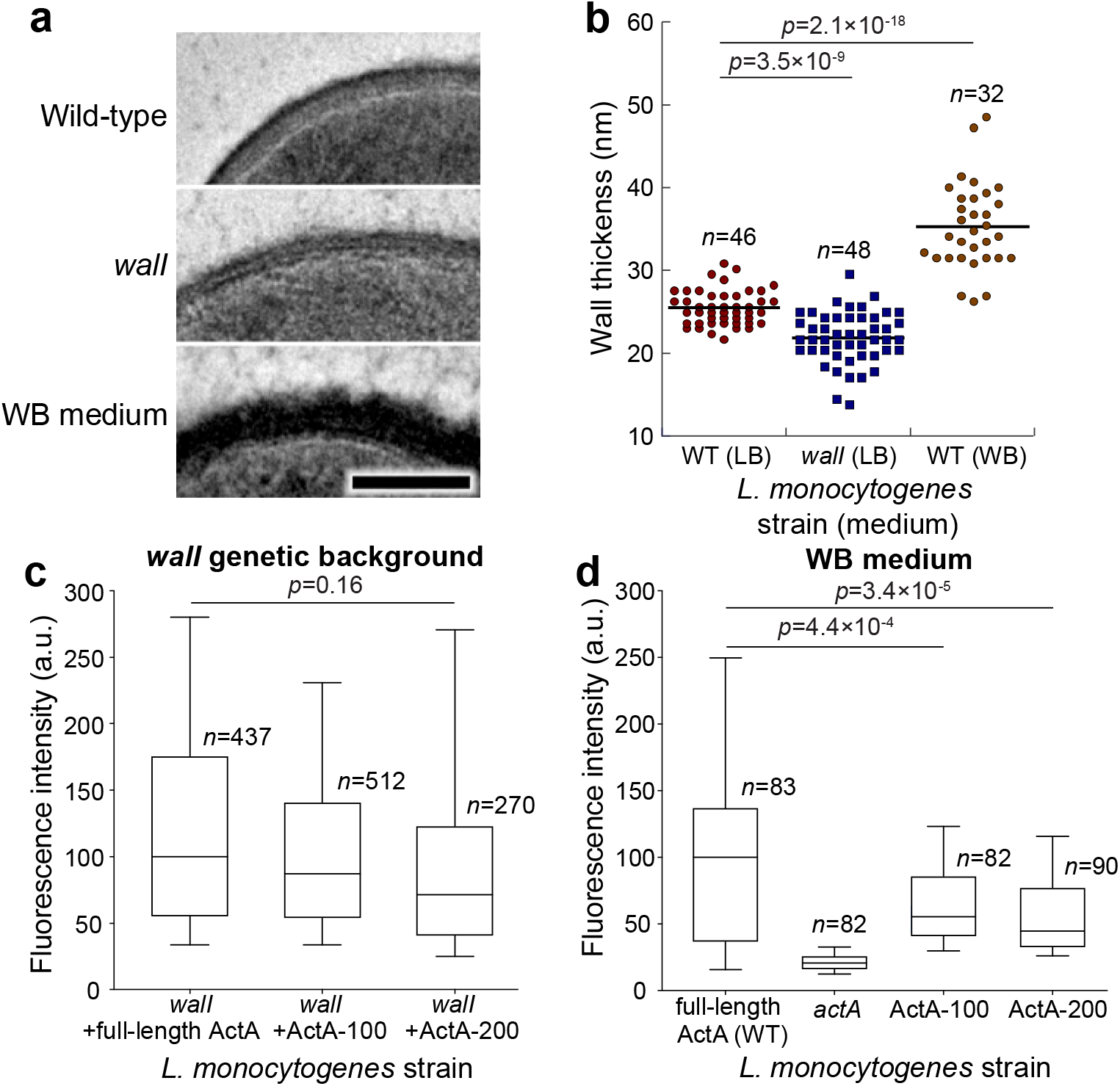
Modulating wall thickness genetically or by changing growth medium affects the critical length for ActA translocation. a) Electron micrographs of wild-type cells in LB medium (top), *walI* cells in LB (middle), and wild-type (WT) cells in WB medium (bottom). b) Deletion of *walI* decreased wall thickness relative to wild-type, and growth in WB increased wall thickness relative to growth in LB. c) In *walI* cells, the thinner cell wall allowed translocation even after truncation of 200 amino acids at the C-terminus. Fluorescence intensities were normalized so that the median of the wild-type strain was 1. The box shows 25^th^ and 75^th^ percentiles with the median as a horizontal line, and whiskers are 10^th^ and 90^th^ percentiles. Blue shading indicates confidence intervals for the median. d) In WB, labeling was significantly lower in both truncation mutants compared with full length ActA. Fluorescence intensities were normalized so that the median of the wild-type strain was 1. The box shows 25^th^ and 75^th^ percentiles with the median as a horizontal line, and whiskers are 10^th^ and 90^th^ percentiles. Blue shading indicates confidence intervals for the median.

Next, we expressed ActA in a strain of *Bacillus subtilis* lacking the major proteases^22^, since *B. subtilis* has a significantly thicker cell wall and a thicker periplasmic space than *L. monocytogenes^11,23^*. Consistent with the predictions of our model, we observed no antibody labeling when expressing full-length ActA from an inducible promoter (Extended Data Fig. 4a). Digesting the cell wall using lysozyme^21^ resulted in ActA labeling (Extended Data Fig. 4b), indicating that the protein was expressed, secreted, and anchored at the bacterial membrane, but was unable to translocate through the *B. subtilis* cell wall. To determine if a longer disordered protein could translocate through the cell wall, we expressed the 1080-amino acid protein iActA from *Listeria ivanovii* in *B. subtilis*. Much like ActA, iActA is a transmembrane protein that promotes actin polymerization at the bacterial surface^24,25^ and is predicted to have an almost entirely disordered N-terminal domain (Extended Data Fig. 4c). When expressing iActA in *B. subtilis*, we observed surface labeling with the antibody (Extended Data Fig. 4d), suggesting that iActA experiences enough confinement within the periplasm to overcome the entropic barrier and navigate across the cell wall.

A compelling prediction of our model is that any sufficiently long disordered protein, otherwise irrespective of its primary sequence, should translocate through a Gram-positive cell wall provided there is sufficient confinement within the periplasm. To test this prediction, we used disordered regions from nuclear pore complex proteins, which were chosen based on having similar lengths and Stokes radii to ActA^26^. To direct these eukaryotic protein regions to the *L. monocytogenes* surface, we created chimeric proteins using the signal sequence and transmembrane domain from ActA; these chimeric proteins were stably produced in *L. monocytogenes* (Extended Data Fig. 5a). Remarkably, the disordered regions from Nsp1 and Nup1 translocated through the cell wall and adopted a polar distribution that was qualitatively similar to ActA localization (Fig. 4a,b and Extended Data Fig. 5b,c). Furthermore, like ActA, Nsp1 polarized in stages and required several bacterial generations to accumulate at poles (Extended Data Fig. 6). Removing the transmembrane domain from the Nsp1 construct resulted in secretion into the extracellular medium and an absence of immunofluorescence signal at the cell surface (Extended Data Fig. 5d), demonstrating that surface labeling required protein anchoring to the membrane, and not any non-specific binding of Nsp1 to the cell wall.

**Figure 4.**
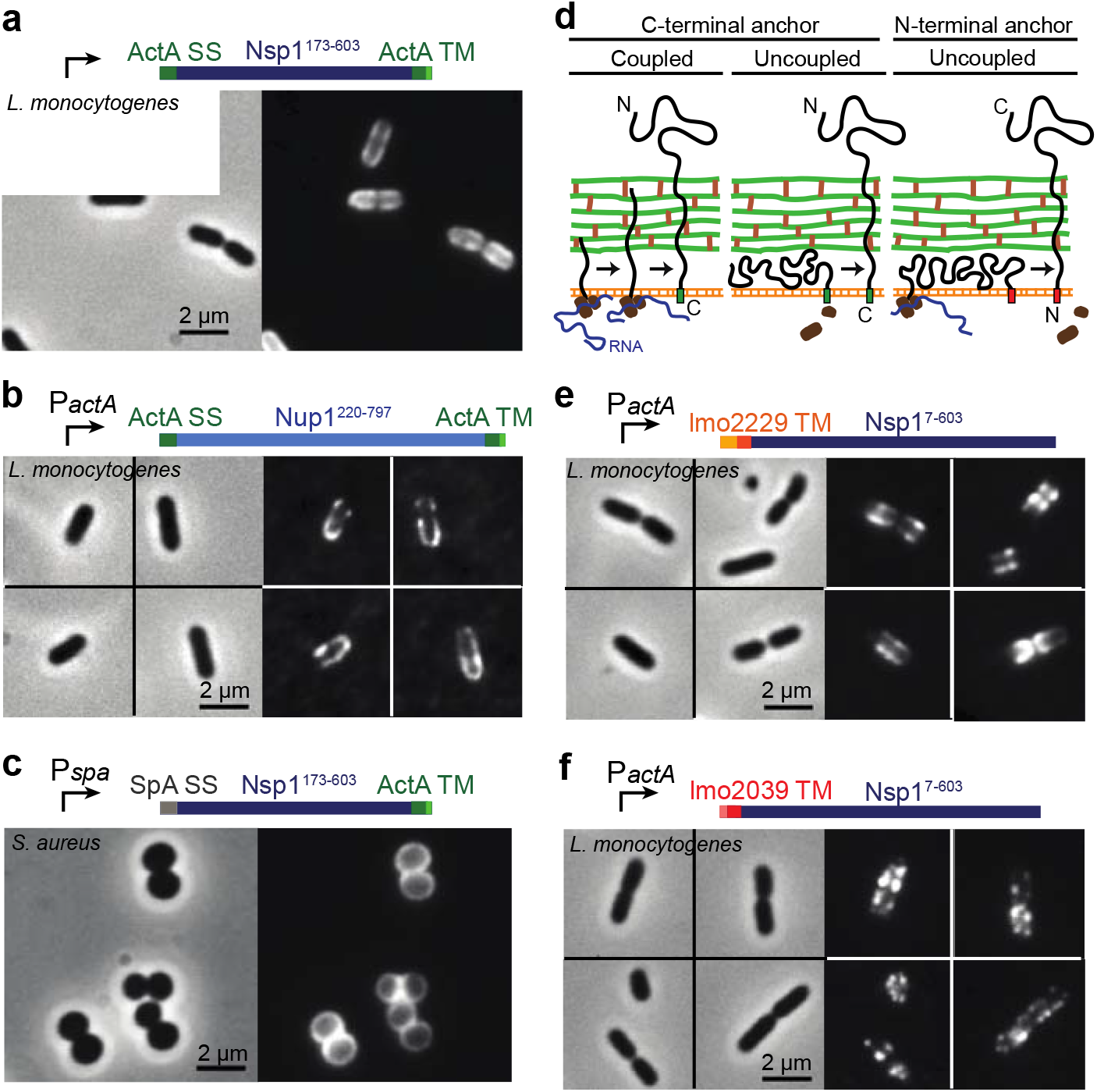
Disordered regions from nuclear pore proteins translocate through the bacterial cell wall, and this process can be uncoupled from secretion through the membrane. Schematics of protein constructs indicating the promoter (*P_actA_* or *P_spa_*), the signal sequence (ActA, Spa, lmo2229, or lmo2039), the protein coding region (Nsp1 or Nup1) and the presence or absence of a C-terminal anchor domain are shown above each set of images, with phase contrast on the left and immunofluorescence on the right. a,b) Immunofluorescence using an antibody against Nsp1, showing surface presentation and polarized localization of (a) Nsp1 and (b) Nup1 in *L. monocytogenes* when expressed with the promoter, signal sequence, and C-terminal anchor from ActA. c) Surface presentation of Nsp1 in *S. aureus*, expressed with the promoter and signal sequence from protein A (SpA). d) Schematic of possible polymer topologies during secretion through the membrane. Secretion and translocation can be either coupled or uncoupled in proteins with a C-terminal anchor (left), while these processes would necessarily be uncoupled for proteins anchored at the N-terminus (right). e) Surface presentation of Nsp1 with an N-terminal transmembrane domain from the PBP lmo2229. Note predominant localization to the mid-cell region. f) Surface presentation and punctate lateral wall localization of Nsp1 with an N-terminal transmembrane domain from the PBP lmo2039.

To further test the generality of our proposed model, we expressed Nsp1 in *Staphylococcus aureus*, a round bacterium with a cell wall thickness comparable to *L. monocytogenes* and a narrower periplasmic thickness than *B. subtilis*^27^. The protein A (SpA) N-terminal signal sequence and ActA C-terminal transmembrane domain were used for Nsp1 secretion and membrane anchoring. We observed antibody-accessible Nsp1 stably presented across most of the *S. aureus* surface (Fig. 4c), similar to the distribution of native protein A^28^, which has been previously shown to be determined by the sequence of the signal peptide^29^. Thus, we have shown that exogenous disordered proteins, even those of eukaryotic origin, can translocate through the cell wall of *L. monocytogenes, B. subtilis*, and *S. aureus*.

Although our model predicts that entropic effects can be sufficient for translocation, we considered the possibility that an active cellular process is also required. For example, the cis-trans-prolyl-isomerase PrsA2 could in principle contribute to ActA translocation as ActA contains several functionally important proline-rich domains. *L. monocytogenes* Δ*prsA2* cells are defective in the secretion of many proteins^30^ including the major secreted hemolysin listeriolysin O (Extended Data Fig. 7a); ActA expression did decrease relative to wild-type in two transposon insertions into *prsA2*, but of the expressed protein the same amount was extractable by SDS as by mechanical disruption, indicating that it was extended beyond the cell wall (Extended Data Fig. 7b). Antibody labeling confirmed translocation of ActA in a *prsA2* mutant (Extended Data Fig. 7c). Moreover, the proline-rich domains of ActA were able to bind the relevant domain of VASP in a *prsA2* mutant (Extended Data Fig. 7d), and *prsA2* bacteria were able to form normal actin-rich comet tails in infected host cells (Extended Data Fig. 7e). Thus, PrsA2 is not required for ActA translocation or for its normal function. We hypothesized that any other active translocation would most likely be a result of mechanical coupling of cell wall translocation to protein secretion through the bacterial membrane, a process that can be driven by the ATPase SecA or the ribosome^18,31^. Coupled force from secretion could possibly push a membrane-anchored protein through the cell wall if the transmembrane anchor were located at the C-terminus (as in native ActA); however, such a secretion mechanism would not be possible for proteins with an N-terminal transmembrane domain, as both ends of the protein would remain associated with the membrane until secretion was complete (Fig. 4d). To test whether mechanical work by the secretion apparatus is required for translocation across the cell wall, we anchored Nsp1 to the membrane using the N-terminal transmembrane domain from *L. monocytogenes* penicillin-binding protein (PBP) lmo2039 or lmo2229. We observed translocation of both lmo2039- and lmo2229-anchored Nsp1 (Fig. 4e,f), indicating that translocation can occur after secretion has completed, with the final orientation of the protein in the opposite state as compared to our previous experiments (C-terminus out, rather than N-terminus out). Constructs containing Nsp1 flanked by both N-terminal and C-terminal transmembrane domains simultaneously failed to translocate across the wall, remaining entirely confined within the periplasmic space (Extended Data Fig. 5e).

To determine whether entropy-driven translocation may be widespread among Gram-positive bacteria, we analyzed 30 bacterial proteomes for surface proteins with long disordered regions. Disordered regions were predicted using the VSL2B predictor^32,33^ and categorized as belonging to a surface protein based on the presence of a predicted transmembrane domain or signal sequence. We found long disordered regions in surface proteins in most Gram-positive species, including disordered regions longer than 500 amino acids in *L. monocytogenes*, *Lactococcus lactis*, and *Lactobacillus*, *Streptococcus* and *Staphylococcus* species (Extended Data Fig. 8 and Extended Data Table 1). Consistent with our model, the *S. aureus* membrane-anchored surface protein EbpS, which extends through the cell wall to bind host elastin^34^, was predicted to have an entirely disordered ectopic domain. In some Gram-positive bacteria, there is a surface structure external to the cell wall, such as an S-layer or mycolate layer, that might negate the entropic benefit of translocation by creating a confined environment beyond the cell wall. For many of the bacterial species that have an S-layer, such as *Clostridium* spp. and *Enterococcus faecalis*, we identified few surface proteins with long disordered regions (Extended Data Fig. 8), suggesting that the need for translocation might drive the evolution of long disordered regions and that entropic translocation is less prevalent or not utilized by these species.

Our results demonstrate that entropy-driven translocation of disordered proteins through a Gram-positive cell wall depends on both the geometric properties of the bacterial cell surface and on the length of the translocating protein, while not requiring any particular amino acid sequence or overall polypeptide orientation. This mechanism is distinct from the entropic process of diffusion down a concentration gradient. While we specifically examined the translocation of disordered membrane-anchored proteins, long disordered regions might also promote the translocation of secreted and cell wall-associated proteins, enabling more efficient navigation through the cell wall. Furthermore, unlike outer membrane biogenesis in Gram-negative bacteria, which requires chaperones^19^ and in some instances is coupled to the hydrolysis of intracellular ATP^35^, this general entropy-driven process appears to be independent of specific biological machinery and energy input associated with secretion.

## Acknowledgments

We thank M. Rexach for the antibody against Nsp1, W. Burkholder, A. Cheung, T. Burke, and D. Portnoy for strains, and J. Lynch, K. Schulz, M. Tsuchida, and S. Weber for comments on the manuscript. Electron microscopy was performed in and with the assistance of the Stanford Cell Sciences Imaging Facility. This work was supported by NIH R37AI-36929 (to J.A.T.), NSF CAREER Award 1149328 (to K.C.H.), NSF grants DBI-0960480, DMS-1616926, and HRD-1547848 (to A.G.) and EF-1038697 (to A.G. and K.C.H.), a James S. McDonnell Foundation Award (to A.G.), the Allen Center for Systems Modeling of Infection (to K.M.N. and K.C.H.), and HHMI (to J.A.T.). The project was supported, in part, by ARRA Award Number 1S10RR026780-01 from the National Center for Research Resources (NCRR); its contents are solely the responsibility of the authors and do not necessarily represent the official views of the NCRR or the National Institutes of Health. K.C.H. is a Chan Zuckerberg Biohub Investigator.

## Author Contributions

D.K.H., F.E.O., K.C.H., and J.A.T. designed the experiments, analyzed the data, and wrote the manuscript. D.K.H., F.E.O., M.J.F., and J.A.T. performed the experiments. D.K.H. and K.M.N. constructed the mutants. A.G. and K.C.H. developed the model. N.S.M. and S.N.M. performed the disorder predictions. All authors reviewed the manuscript before submission.

## Author Information

The authors declare no competing financial interests.

## Extended Data Figures

**Extended Data Figure 1:**
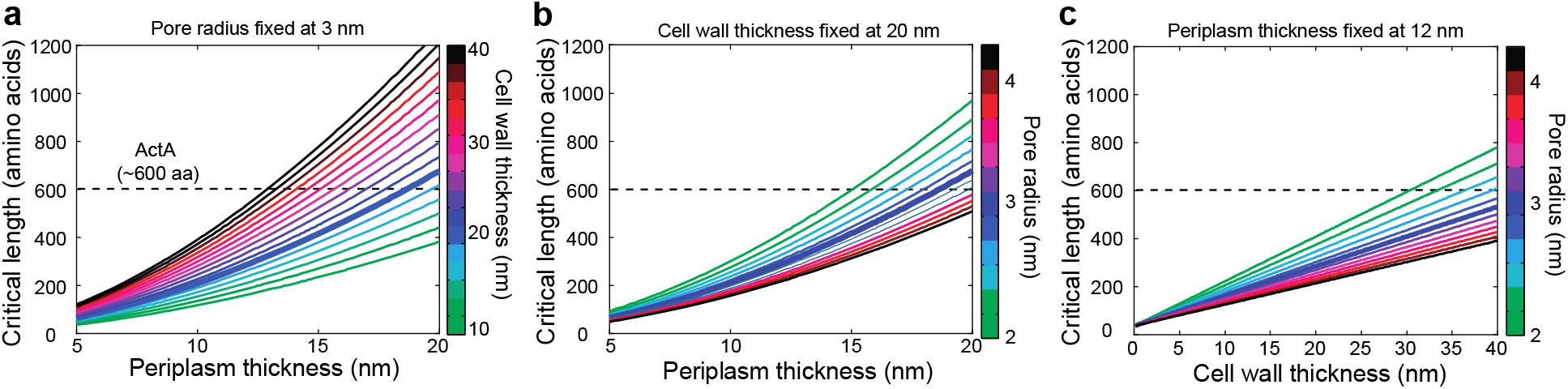
Predicted critical lengths for an ActA-like polymer for different values of cell wall thickness, periplasm thickness and pore radius. a) Critical length as a function of periplasm thickness *P* for different values of cell wall thickness *w*, with a pore radius *R* set at 3 nm. Each curve denotes a different cell wall thickness, with *w* ranging from 10 nm to 40 nm. b) Critical length as a function of periplasm thickness *P* for different values of *R*, with *w* set at 20 nm. c) Critical length as a function of cell wall thickness *w* for different values of *R*, with *P* set at 12 nm. In (b,c), each curve denotes a different pore radius, with *R* ranging from 2 nm to 4.2 nm. For all graphs the bold line signifies parameters used in Fig. 1: *w* = 20 nm, *P* = 12 nm, *R* = 3 nm. The dashed horizontal line denotes the length of an ActA molecule. The entropic model predicts that any parameter set resulting in a critical length falling below this dashed line will result in proper translocation of ActA. All parameter values were chosen so that 2*R* and *P* are substantially more than the persistence length of ActA (Supplementary Text).

**Extended Data Figure 2:**
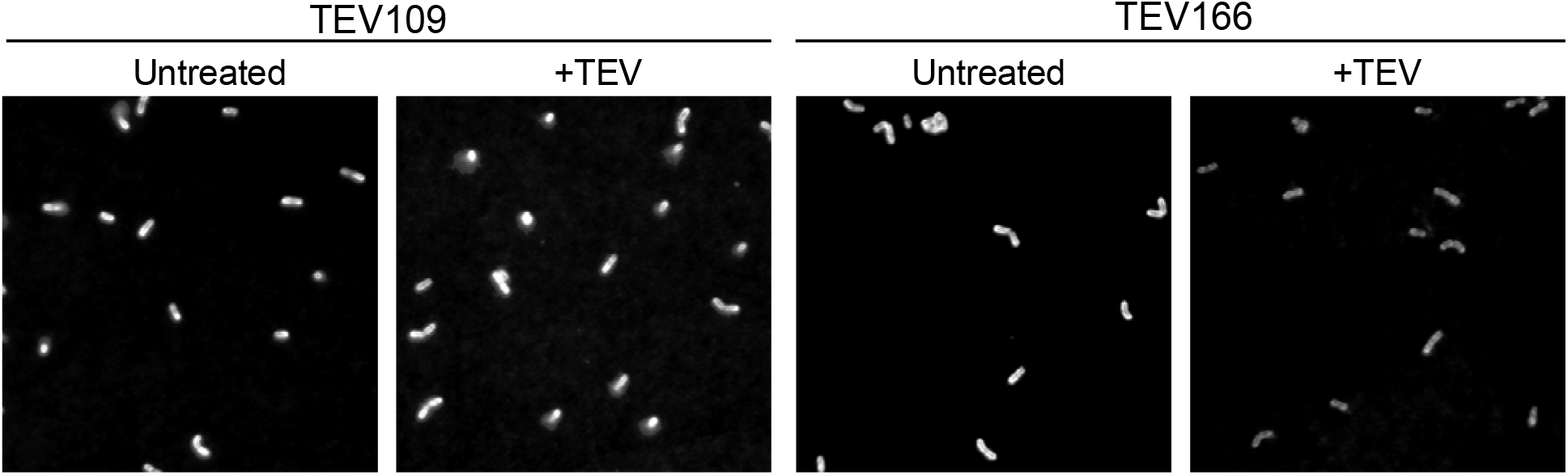
Effects of TEV protease treatment on surface presentation of ActA. A representative TEV cleavage experiment showing population distributions of untreated and TEV protease-treated conditions for strains in Fig. 2d,e.

**Extended Data Figure 3:**
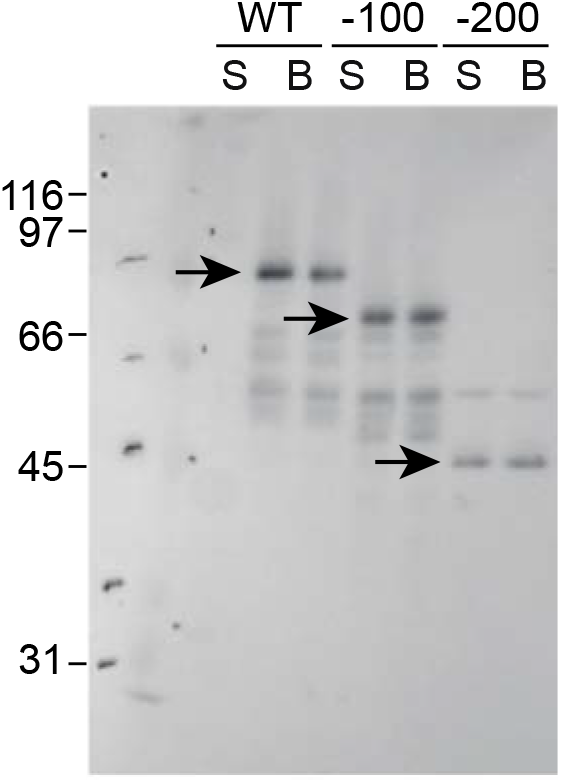
Genetic truncations of ActA were detected at expected lengths. −100 and −200 refer to 100- and 200-amino acid truncation mutants, respectively. SDS treatment (S) and mechanical disruption by bead-beating (B) as in Fig. 1a,b were performed for all strains. Arrows indicate (from left to right) full length ActA, the 100-amino acid truncation, and the 200-amino acid truncation.

**Extended Data Figure 4:**
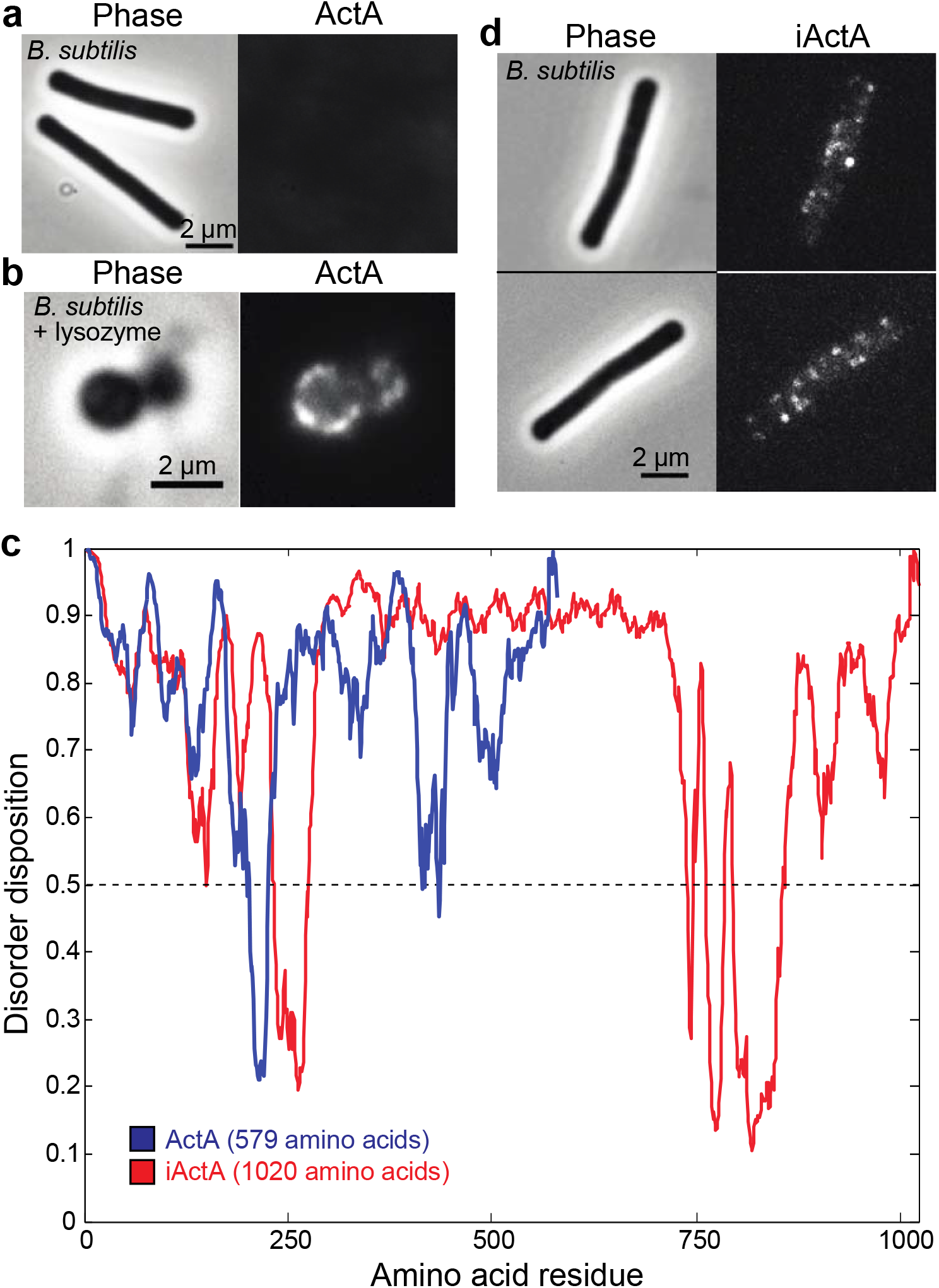
The disordered transmembrane protein iActA extends through the *B. subtilis* cell wall. a) *B. subtilis* cells expressing the 65-kDa ActA protein exhibited no appreciable immunofluorescence labeling, indicating minimal translocation. b) Labeling of lysozyme-treated *B. subtilis* expressing ActA showed ActA associated with the membrane of the spheroplast. c) Disorder prediction results for the extracellular regions of ActA and iActA using the PONDR-FIT predictor. d) Punctate immunofluorescence labeling was observed in *B. subtilis* engineered to express the 108-kDa iActA protein, suggesting that a longer disordered protein could translocate across the relatively thick *B. subtilis* cell wall compared with that of *L. monocytogenes*.

**Extended Data Figure 5:**
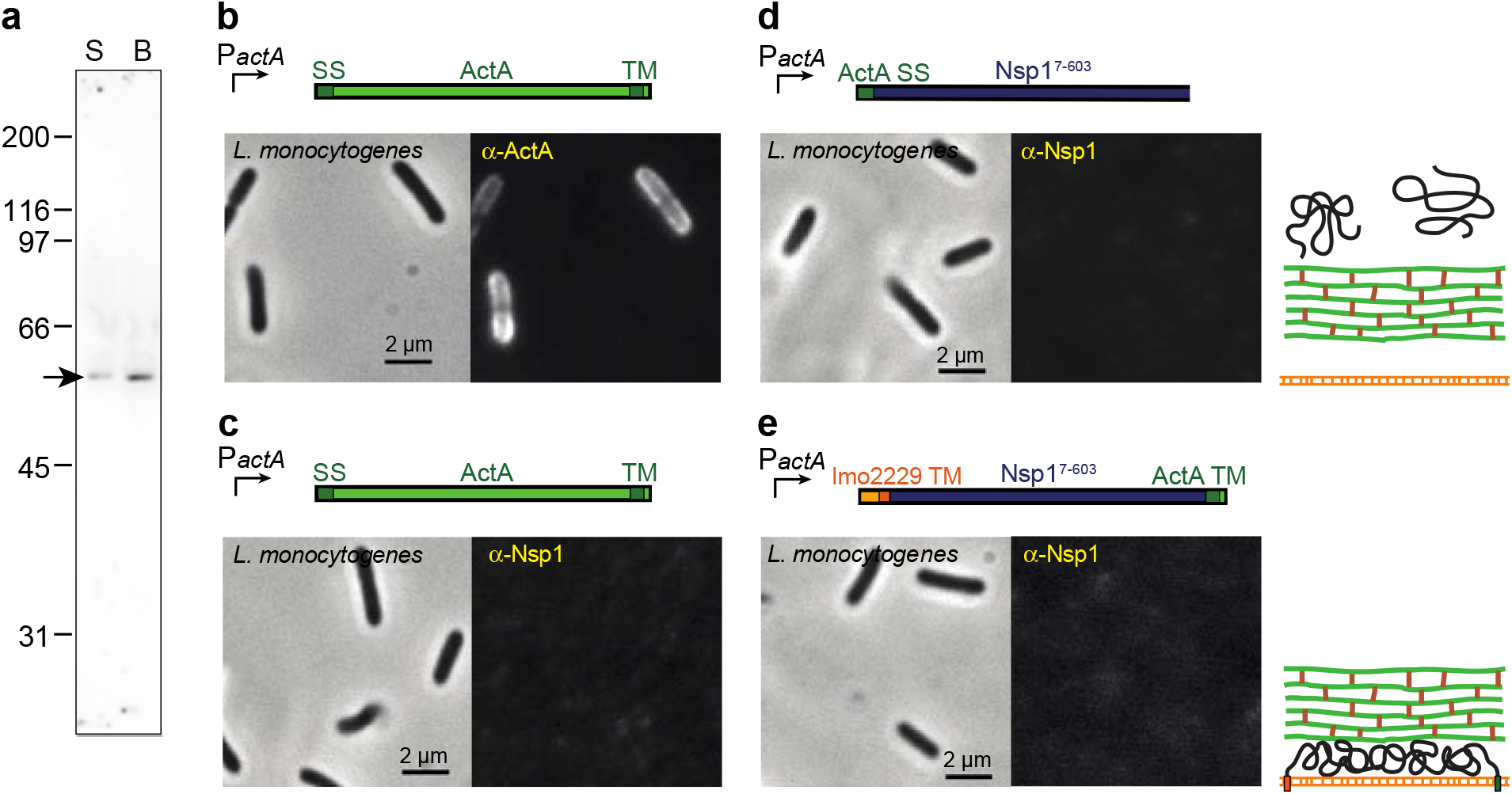
Lack of immunofluorescence signal for Nsp1 constructs that are secreted or anchored by transmembrane domains at both termini. a) Western blotting showed stable expression of the Nsp1 chimeric construct (arrow) from Fig. 4a,b. b) Polar distribution of ActA in the wild-type strain of *L. monocytogenes* using an antibody against ActA. c) Lack of signal in the wild-type strain using an antibody against Nsp1. d) Immunofluorescence of Nsp1 expressed with an amino-terminal signal sequence from ActA but no transmembrane anchor. e) Nsp1 expressed with an amino-terminal transmembrane domain from lmo2229 and carboxy-terminal5 transmembrane domain from ActA. No visible labeling was observed when Nsp1 was anchored at both ends, confirming that antibodies do not label proteins that are trapped in the periplasm.

**Extended Data Figure 6:**
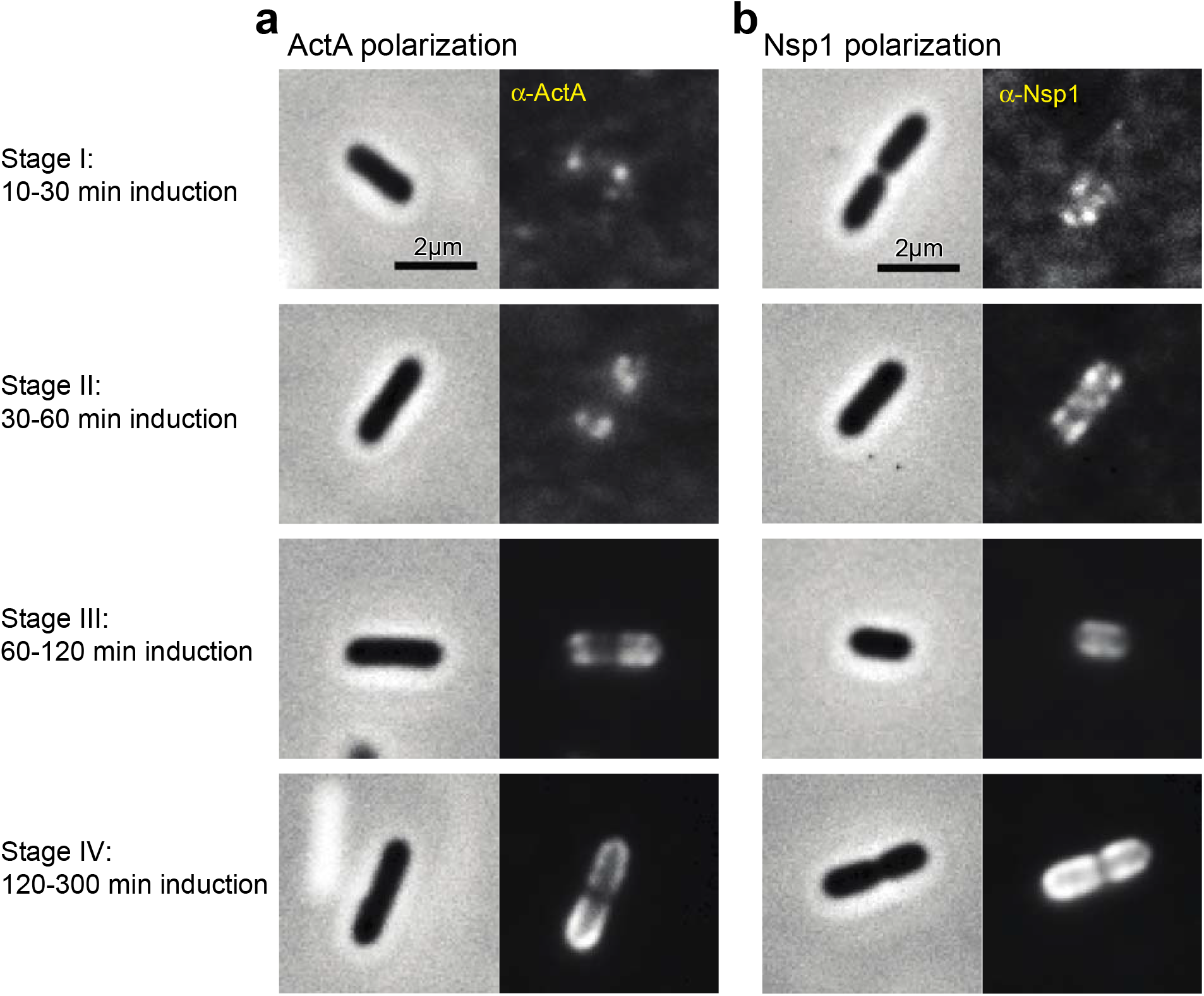
Time course of disordered protein localization and polarization in *L. monocytogenes*. a) ActA expressed from the ActA promoter. b) Nsp1, with a signal sequence and transmembrane domain from ActA, expressed from the ActA promoter.

**Extended Data Figure 7:**
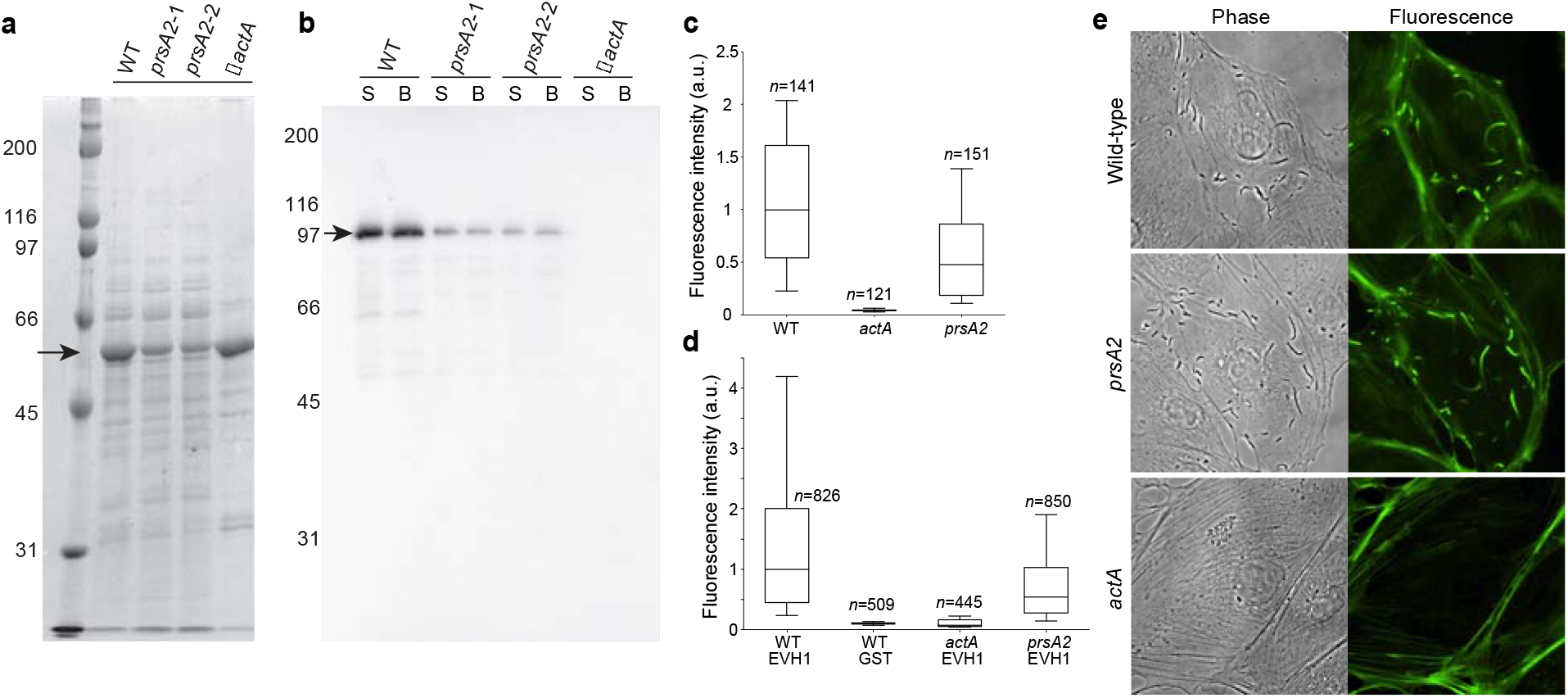
ActA expression, translocation, and exposure, as well as VASP binding and comet-tail formation, were maintained in *ΔprsA2* cells. a) Expression of listeriolysin O (arrow) was lower in two *prsA2* transposon insertion mutants, as expected, but similar to in a Δ*actA* mutant, as shown in a Coomassie-stained gel of TCA-precipitated culture supernatant. b) Expression of ActA (arrow) in *prsA2* cells decreased to a similar extent as listeriolysin O in (**a**); no expression was detected in Δ*actA* cells. Levels were similar between cells treated with SDS (S) or subjected to bead beating (B), indicating that most ActA was translocated across the cell wall. c) Immunofluorescence labeling confirmed that ActA was exposed on the surface of wild-type and*prsA2* cells. d) The EVH1 domain of VASP fused to GST bound to the surface of *prsA2* and wild-type cells expressing wild-type ActA, but not to a Δ*actA* cells. GST alone also did not bind the surface of wild-type bacteria. Shown is immunofluorescence labeling of GST. e) Wild-type and *prsA2 L. monocytogenes* formed comet tails in host cells, but Δ*actA L*. *monocytogenes* did not. Thus, PrsA2 is not required for ActA translocation or function.

**Extended Data Figure 8:**
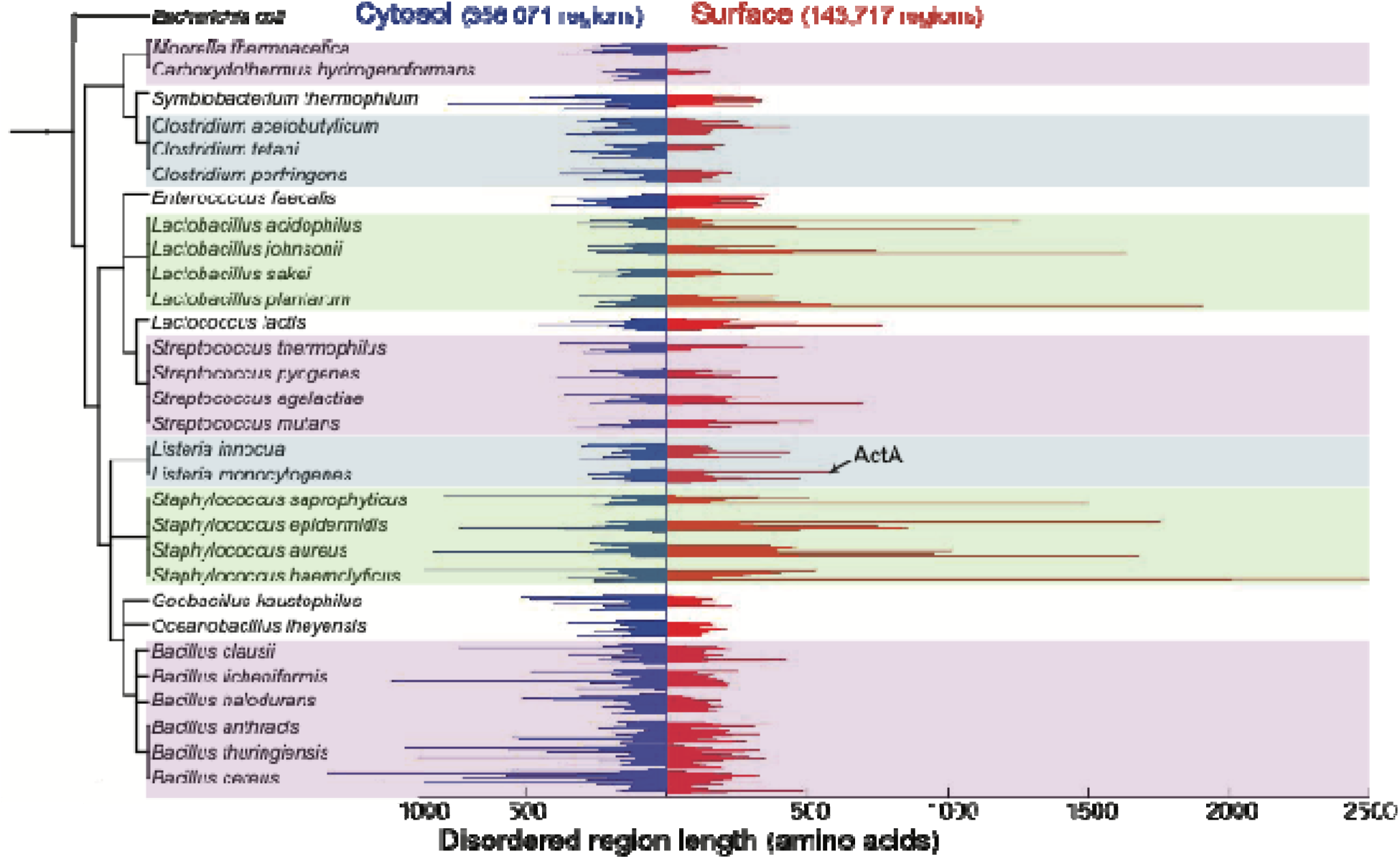
Prevalence of long disordered regions in surface proteins of Gram-positive bacteria. Surface proteins (disordered regions denoted as red bars) were identified based on the presence of a predicted transmembrane domain, signal sequence, or LPXTG motif. Proteins that were not predicted to be surface proteins were considered cytosolic (disordered regions denoted as blue bars). Disordered regions were determined using the disorder predictor VSL2B^32,33^. The shaded regions highlight groups of phylogenetically related bacteria. See also Extended Data Table 1.

**Extended Data Table 1.** List of surface proteins with a predicted disordered region of at least 250 amino acids for *L. monocytogenes, S. aureus, L. acidophilus, E. faecalis, S. pyogenes, B. anthracis*, and *L. lactis*.

| Species | Gene | Function | Disordered region length |
| --- | --- | --- | --- |
| *Listeria monocytogenes* | *actA* | actin-assembly inducing protein | 579 |
|  | *lmo1105* | hypothetical protein | 273 |
|  | *lmo1799* | Peptidoglycan-binding protein | 473 |
|  | *lmo2287* | phage tape-measure protein | 283 |
| *Staphylococcus aureus* | *spa* | IgG-binding protein A | 368 |
|  | *coa* | staphylocoagulase | 438 |
|  | *geh* | Glycerol-ester hydrolase | 265 |
|  | *sdrC* | Ser-Asp-rich fibrinogen-binding, bone sialoprotein-binding protein | 285 |
|  | *sdrD* | Ser-Asp-rich fibrinogen-binding, bone sialoprotein-binding protein | 336 |
|  | *clfA* | fibrinogen-binding protein A, clumping factor | 461 |
|  | *SA0743* | hypothetical protein, similar to staphylocoagulase | 295 |
|  | *htrA* | serine protease | 392 |
|  | *ebhA* | hypothetical protein,<br>similar to streptococcal<br>adhesin Emb | 383; 517; 260; 457; 1009;<br>739; 369; 821; 429 |
|  | *ebhB* | hypothetical protein,<br>similar to streptococcal<br>adhesin Emb | 921 |
|  | *ebpS* | elastin-binding protein | 316 |
|  | *SA1559* | hypothetical protein,<br>similar to caldesmon | 398 |
|  | *SA1577* | hypothetical protein,<br>similar to FmtB | 301 |
|  | *fmtB* | FmtB protein | 948; 418 |
|  | *fnbB* | fibronectin-binding protein | 468 |
|  | *fnb* | fibronectin-binding protein | 469 |
|  | *clfB* | clumping factor B | 324 |
|  | *SA2439* | hypothetical protein | 386 |
|  | *SA2447* | hypothetical protein,<br>serine-rich adhesin for<br>platelets | 1673 |
|  | *lip* | triacylglycerol lipase | 265 |
| *Lactobacillus*<br>*acidophilus* | *LBA1020* | mucus-binding protein | 256 |
|  | *LBA1019* | mucus-binding protein | 260 |
|  | *fmtB* | surface protein | 1251; 1098 |
|  | *LBA1637* | membrane protein | 461 |
|  | *LBA1633* | surface protein | 591 |
|  | *LBA1634* | surface protein | 376 |
| *Enterococcus faecalis* | *EF0093* | cell wall surface anchor<br>family | 359 |
|  | *EF0146* | surface exclusion protein | 349; 341 |
|  | *EF0502* | hypothetical protein | 314 |
|  | *EF1099* | collagen adhesin protein | 344 |
|  | *EF1523* | hypothetical protein | 316 |
|  | *EF1877* | hypothetical protein | 283 |
|  | *EF2318* | M24/M37 family peptidase | 327 |
|  | *EF3023* | polysaccharide lyase<br>family | 338 |
|  | *EF3173* | hypothetical protein | 308 |
|  | *EF3314* | cell wall surface anchor<br>family | 289 |
| *Streptococcus pyogenes* | *spyM18_0256* | surface exclusion protein | 266 |
|  | *spa* | protective antigen | 394 |
|  | *emm18* | M18 protein | 339 |
| *Bacillus anthracis* | *BAS0454* | phage tape-measure protein | 371 |
|  | *BAS1009* | hypothetical protein | 316 |
|  | *BAS3021* | cell wall anchor domain<br>protein | 331 |
|  | *BAS4630* | spore germination protein | 283 |
| *Lactococcus lactis* | *lcnD* | lactococcin A ABC<br>transporter permease | 260 |
|  | *yihD* | hypothetical protein | 459 |
|  | *yjaE* | hypothetical protein | 276 |
|  | *yoiC* | hypothetical protein | 509 |
|  | *yqfG* | hypothetical protein | 767 |
|  | *usp45* | hypothetical protein | 316 |

## Supplementary Text

### Model

Previous theoretical work has investigated polymer translocation through a porous entropic barrier^36–43^. Here, we present a model for the translocation of an intrinsically disordered protein through the Gram-positive cell wall, with the goal of making qualitative predictions for how the critical length should scale with the key variables of cell-wall pore size and periplasmic/cell wall thickness; the critical length will depend on these variables (Extended Data Fig. 1) but the qualitative conclusions of our study are general features of the model.

#### Cell surface properties

The bacterial cell surface was modeled using three parameters: cell wall thickness (*w*), periplasmic thickness (*P*), and cell-wall pore radius (*R*). To estimate cell wall thickness, we used previous measurements from conventional and cryogenic transmission electron microscopy (cryo-TEM)^11,23,27,44,45^ as well as our own measurements (Fig. 3b). Cryo-TEM measurements were used to approximate the periplasmic thickness^11,23,27^; a distinct periplasmic space in *L. monocytogenes* has not been observed using conventional TEM, possibly because of poor preservation of the structure during fixation, dehydration, and/or embedding. For pore size, we modeled a continuous pore through the cell wall. While a continuous pore has not been observed, the cross-linked nature of the sacculus creates an inherently porous structure, and the quantitative predictions of our model do not depend on the precise geometry of the pore. Isolated sacculi of *Escherichia coli* and *Bacillus subtilis* have a mean pore radius of ~2-3 nm as measured by permeation of fluorescent dextrans^8^, suggesting a typical pore size that may be a molecular property of peptidoglycan and hence reasonably conserved across bacterial species.

#### Kuhn length of ActA

The extracellular region of ActA has no apparent secondary structure and has a Stokes radius *R_H_* of ~8 nm^16^. Because of the lack of secondary structure, we used a simple polymer physics model to describe the behavior of ActA^46^. For a self-avoiding polymer, the radius of gyration *R_G_* can be estimated as 1.58*R_H_*^47^. The estimated persistence length of such a polymer is

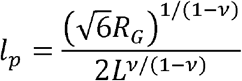

where *L* is the length of the completely stretched protein and *v* is the Flory exponent. For a freely jointed chain, *v*=1/2 and 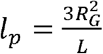, but for more realistic scenarios, *v*>0.5^48^. For the plots in Fig. 1, 2, and Extended Data Fig. 1, we model ActA as a self-avoiding polymer with *v*=0.59^42^; the qualitative predictions of our model are insensitive to the choice of *v*. For a protein of 600 amino acids, the completely stretched length would be approximately 228 nm, assuming each amino acid occupies 0.38 nm of length. Thus, the persistence length of ActA is *l_p_* = 0.875 nm, giving an effective Kuhn length of *a* = 2*l_p_* ≈ 1.75 nm or 4.61 amino acids. Hereafter, we refer to all lengths in units of *a*.

#### Blob approximation

A polymer subject to confinement or tension has a distribution of conformations whose statistical properties can be approximated as those of a connected series of “blobs”, wherein each blob represents the maximal size of a segment of the polymer below which the confinement and tension effects can be ignored^42^. Scaling then argues that the number of subunits in each blob of size *r* is *g□r*^1/*v*^ (assuming *g*>> 1 or *r>l_p_*, which is the case for the typical confinements and extensions we consider).

#### Entropic cost of confinement

To estimate the entropy of a polymer confined within a periplasm of thickness *P*=2*p*, we assume that *n_p_»p*, where *n_p_* is the number of Kuhn lengths of the polymer situated in the periplasm. The number of effective blobs in the periplasm is *N_b_* = *n_p_/g*, with *g* ~ *p*^1/*v*^. The entropic cost *F* of polymer confinement in the periplasm scales as *k_B_T*per blob^42^, giving

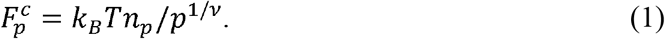

Using a similar argument^42^, the entropic cost of confinement of a polymer in a pore of radius *R* in the cell wall scales as 1/*R*^1/*v*^.

#### Entropic cost of stretching

A polymer will also incur an entropic cost if it is stretched beyond its preferred, unperturbed size. If a polymer of unperturbed radius less than *p* is stretched across the periplasm, it can be regarded as a chain of blobs of radius *r*. Each blob will have *g* subunits, where *g* ~ *r*^1/*v*^, and the number of blobs needed to cross the periplasm is *p*/*r*. Thus, the total number of subunits in the periplasm is

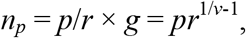

which results in the relation *r* = (*n_p_*/*p*)^*v* /(1-*v*)^. Thus, the stretching free energy cost is^42^

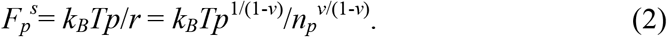

A similar argument can be made for stretching across the cell wall, replacing *p* with *w*.

#### Free energy cost

To determine the free energy landscape, we calculated the free energy cost of each configuration of the polymer relative to a polymer of the same length in an unconfined solution as a function of *p*, *w*, *R*, and *n*. Using Eq. 1 and 2, the free energy costs in the periplasm and cell wall are

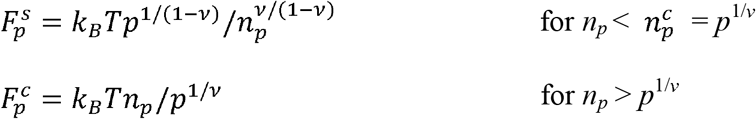

and

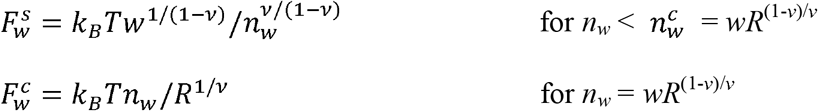

respectively. We do not consider *n_w_* > *wR*^(1-*v*)/*v*^ (at which point the blobs are of size *R*) as any additional polymer subunits can be moved to the unconfined exterior or the periplasm where there is less confinement and therefore results in lower free energy.

#### Critical length

To calculate the critical length for polymer translocation, we determined the minimum free energy cost *E*_cross_ for crossing the periplasm and cell wall, and compared this value with the total free energy cost for the entire polymer to be confined in the periplasm. Translocation through the cell wall becomes favorable when the free energy cost of periplasmic confinement *E*_peri_ is greater than *E*_cross_, although there is an energy barrier for cell wall crossing. The condition that determines the critical length *n_c_* is when *E*_peri_ = *E*_cross_:

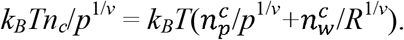

The critical length is then given by

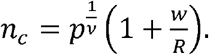

#### Fraction of molecules extending through the cell wall

The expected fraction of molecules in an extended state at equilibrium is

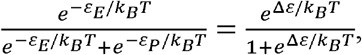

where *ε_E_* and *ε_P_* are the free energy costs for a polymer in an extended state and periplasmic state, respectively, and Δ*ε* = *ε_P_-ε_E_*.

## Online Methods

### Strains and growth conditions

Unless otherwise noted, all strains were grown with aeration at 37 °C.

### *Listeria monocytogenes* growth conditions

TEV-cleavage experiments were performed using a PrfA* strain of *L. monocytogenes* that constitutively expresses ActA^49^. These strains were grown for 12-16 h in 1.5X LB medium containing 7.5 μg mL^-1^ chloramphenicol (Cm; Sigma) until reaching an OD_600_=0.4-0.5. All other experiments were performed in the 10403S background. *actA* expression (10403S strains) from the native promoter^4^ or an IPTG-inducible promoter^50^ was induced as previously described. Bacteria were grown overnight in BHI medium containing 7.5 μg mL^-1^ Cm, and then overnight cultures were spun down and resuspended in the appropriate medium. To express ActA from the SPAC/*lac*Oid promoter, cultures were diluted 1:50 into BHI medium containing 7.5 μg mL^-1^ Cm and 0.5 mM IPTG. To express ActA from its native promoter, cultures were diluted 1:25 into an induction medium of LB supplemented with 25 mM α-D-glucose 1-phosphate (Sigma) and ~1-2% (w/v) activated carbon pellets^51,52^.

### *Bacillus subtilis* growth conditions

Strain WB600^22^ was grown overnight in LB medium containing 5 μg mL^-1^ Cm, and then diluted 1:25 into LB medium containing 5 μg mL^-1^ Cm and grown until early-log phase (OD_600_=0.2-0.25). Cultures were then spun down and resuspended in an equal volume of LB medium containing 5 μg mL^-1^ Cm (pH 6.0) and grown for 2 h. ActA and iActA were expressed from the *gsiB* promoter, which is induced during acid stress^53,54^.

### *Staphylococcus aureus* growth conditions

Strain RN6390 (Δ*spA*) was grown overnight in TSB medium containing 5 μg mL^-1^ Cm, and then diluted 1:50 into TSB medium containing 5 μg mL^-1^ Cm and grown until mid-log phase (OD_600_=0.5-0.7). For *S. aureus*, Nsp1 expression was driven by the *spA* promoter.

### Strain construction

For *L. monocytogenes* constructs, the site-specific bacteriophage integration vectors pPL1 and pPL2 were used as previously described^55^. pPL1 was used for ActA constructs with an internal TEV recognition site, while pPL2 was used for nuclear pore complex protein constructs. For IPTG-inducible expression of ActA, we used the previously described *L. monocytogenes* expression vector pHLIV2^50^, which is an integration vector derived from pPL2 (strain created by Michelle Rengarajan, Theriot laboratory). For expression of ActA and iActA in *B. subtilis*, we used the previously described expression vector pHCMC03^54^. We used the shuttle vector pOS1^56^ for Nsp1 expression in *S. aureus*. Disruption of *walI* in ActA truncation mutants was performed by U153 phage transduction of a *walI::Tn* mutant (obtained from the Portnoy laboratory) into the appropriate truncation mutant strain.

Overlap extension polymerase chain reaction was used to insert the TEV protease recognition site into ActA. The DNA sequence of the TEV recognition site was used as the overlapping region. Desired sequences, as specified in the **Strain Information** section, were amplified from genomic preps of *L. monocytogenes* strain 10403S (ActA; lmo2229; lmo2039), *L. ivanovii* strain ATCC19119 (iActA), *Saccharomyces cerevisiae* strain S288C (Nsp1; Nup1), or *S. aureus* strain RN4220 (SpA).

### Slide preparation

18-mm #1.5 coverslips were cleaned in concentrated HCl for 4 h, followed by extensive washing with ddH_2_O. Coverslips were then rinsed in 95% ethanol, and then in 100% ethanol before being transferred to a glass petri dish and autoclaved.

### Immunofluorescence

Unless otherwise noted, bacteria were fixed using formaldehyde at a final concentration of 3.7%. Sixty microliters of fixed bacterial culture was placed on an acid-washed coverslip and incubated at room temperature for 10-30 min depending on culture density. Coverslips were washed twice with PBS, and then incubated with primary antibody at the appropriate dilution for 1 h. Antibodies were used at the following dilutions: rabbit polyclonal anti-peptide antibody against the first proline-rich repeat of ActA, 1:500; rabbit polyclonal antibody against full length ActA, 1:2000; rabbit polyclonal antibody against the FG-region of Nsp1 (generous gift from Michael Rexach, University of California at Santa Cruz) or Abcam anti-Nsp1 antibody 32D6, 1:2000. Coverslips were washed twice with PBS, and then incubated with Alexa Fluor-488 goat anti-rabbit antibody (Invitrogen) at 1:1000 for 1 h. Coverslips were washed three times with PBS, blotted to remove excess liquid, and sealed using VALAP (vaseline:lanolin:paraffin; 1:1:1).

For TEV cleavage experiments, we used a modified protocol to maximize the number of bacteria lying flat on the coverslip: bacteria that are only partially attached to the coverslip often have regions that drift out of the focal plane and therefore cannot be used for analysis. For these experiments, 60 μL of bacterial culture were placed on a 22-mm #1.5 polyL-lysine-coated coverslip. After a 5-min incubation at room temperature, coverslips were blotted, and 50 μL of 3% glutaraldehyde were added to fix the bacteria to the coverslip. After 10 min, coverslips were washed twice in PBS and immunofluorescence was performed using the proline-rich repeat antibody against ActA, as described above.

### Microscopy

All imaging was performed on an upright Zeiss Axioplan 2 microscope equipped with a 100X 1.4 NA Plan Apo objective lens and a 1024×1024-pixel back-illuminated EMCCD camera (Andor) using MetaMorph software.

### Image analysis

Morphometrics^57^ was used to analyze images from the TEV cleavage experiments. Briefly, segmentation of phase contrast images was used to determine cell outlines. Next, the mean fluorescence intensity for each cell was calculated as the total intensity within a cell outline minus the mean background fluorescence, divided by the area of the cell. Normalization was performed by dividing the mean fluorescence intensity of a cell by the average intensity per cell for the untreated control population. Five independent TEV experiments were performed.

### Electron microscopy

Bacterial strains were grown overnight for 16 h in the stated medium at 37 °C. Fixation and processing was performed using conventional procedures for TEM on bacterial samples. Briefly, ~2 mL of culture was spun down and fixed in 50 μL of 2% glutaraldehyde and 4% paraformaldehyde in 0.1 M sodium cacodylate buffer (pH 7.3) for 1 h at room temperature, followed by post-fixation in 1% osmium tetroxide for 1 h at 4 °C and then 1% uranyl acetate overnight. Dehydration through ethanol was followed by acetonitrile and finally embedding in Embed 812 (EMS #14120). Imaging of thin sections was performed on a JEOL1400 TEM operating at 120 kV and 10,000X with a Gatan Orius 10.7 megapixel CCD camera. All steps from post-fixation to sectioning were conducted by staff at the Stanford Cell Sciences Imaging Facility. To measure cell wall thickness, images showing a section through the midplane of a bacterial cell were chosen and distances were quantified using ImageJ.

### TEV cleavage

For each strain, 12 μL of TEV buffer (1 M Tris-HCl pH 8; 10 mM EDTA) and 2 μL 100 mM DTT were added to 285 μL of growing culture. One hundred microliters of this mixture were aliquoted into two Eppendorf tubes, and 1 μL of purified TEV protease (1.1 mg mL^-1^; purified from *E. coli* expressing His-tagged S219V TEV protease using plasmid pRK793 (Addgene)) was added to one of the tubes. Tubes were incubated at 30 °C for 45 min. After 45 min, cells were washed and then 60 μL were used for immunofluorescence, while the remaining 40 μL were used for Western blot analysis.

### Western blotting

To extract ActA, bacteria were boiled in SDS-PAGE loading buffer, as previously described^16^. The bacterial cell wall was digested prior to sample preparation by adding *Listeria*-specific phage lysin Ply118^58^. The primary antibody against the proline-rich repeat of ActA was used at a 1:10,000 dilution. Detection was with a chemiluminescent detection system using a horseradish peroxidase (HRP)-conjugated goat anti-rabbit secondary antibody at 1:20,000 (SouthernBiotech)^4^.

### SDS PAGE

SDS PAGE was performed as previously described^59^. SDS samples of bacteria were made to 2.3 OD_600_ units mL^-1^ in sample buffer. For culture supernatants, the equivalent of 1 OD_600_ of bacterial culture was clarified by centrifugation and the remaining proteins concentrated to 150 μL in sample buffer by DOC/TCA precipitation as previously described^60^.

### Predicting disordered regions and surface proteins

Disordered regions for each protein from 294 prokaryotic species have previously been predicted^32^ using the VSL2B predictor^33^. Proteins were categorized as surface proteins based on the presence of a predicted transmembrane domain by TMHMM v2.0^61^, signal sequence by SignalP v3.0^62^, or LPXTG motif^63,64^. Using these predictions, every disordered region of each protein was plotted as a cytosolic or surface protein (Extended Data Fig. 8). The phylogenetic tree was generated using Interactive Tree Of Life^65^. In Extended Data Table 1, surface proteins with disordered regions longer than 250 amino acids are listed for selected bacterial species.

### Statistical analyses

The significance of differences between two distributions was computed using a two-sample t-test.

### Data availability

All data are available upon request from Julie Theriot.

### Code availability

All custom scripts are available upon request from Kerwyn Casey Huang.

## Additional strain information

**JAT817** [*PgsiB-actA*^1-639^]

*PgsiB: gsiB* promoter, activated in response to a variety of stresses. ActA^1-639^: full length ActA from *L. monocytogenes*.

**JAT818** [*PgsiB-iactA*^1-1080^]

iActA^1-1080^: full length iActA from *L. ivanovii*.

**JAT836** [*tRNAA^rg^::PactA-actA*^1-30^ *-nup1*^220-797^-*actA*^600-639^]

*PactA: actA* promoter. *ActA*^1-30^ : signal sequence of ActA. Nup1^220-797^ : previously characterized disordered region of the Nup1 nuclear pore protein^23^. ActA^600-639^ : transmembrane region of ActA, which includes a short cytosolic tail of a few amino acids. This construct has a carboxy-terminal transmembrane domain.

**JAT840** [*tRNA^Arg^::PactA-actA*^1-30^-*nsp1*^173-603^-*actA*^600-639^]

*nsp1*^173-603^ : previously characterized disordered region of the Nsp1 nuclear pore protein^23^. This construct has a carboxy-terminal transmembrane domain.

**JAT889** [*tRNA^Arg^::PactA-lmo2229*^1-61^-*nsp1*^7-603^]

lmo2229^1-61^: transmembrane region of lmo2229, which includes a cytosolic tail of around 40 amino acids. Nsp1^7-603^: disordered region of the Nsp1 nuclear pore protein. This construct has an amino-terminal transmembrane domain.

**JAT890** [*tRNA^Arg^::PactA-lmo2039*^1-31^-*nsp1*^7-603^]

lmo2039^1-31^: transmembrane region of lmo2039, which includes a cytosolic tail of around 10 amino acids. This construct has an amino-terminal transmembrane domain.

**JAT915** [*tRNA^Arg^::PactA-actA*^1-30^-*nsp1*^7-603^]

Nsp1^7-603^ with an amino-terminal signal sequence (ActA^1-30^) but no transmembrane domain.

**JAT1006** [*tRNAA^rg^::PactA-lmo2229*^1-61^-*nsp1*^7-603^-*actA*^600-639^]

Nsp1^7-603^ with amino-terminal (lmo2229^1-61^) and carboxy-terminal (ActA^600-639^) transmembrane domains.

**JAT1024** [*comK::PactA-actA*^1-6^-*TEV-actA*^64-639^]

ActA with TEV sequence placed between amino acids 63 and 64 (relative to start codon).

**JAT1025** [*comK::PactA-actA*^1-139^-*TEV-actA*^140-639^]

ActA with TEV sequence placed between amino acids 139 and 140.

**JAT1026** [*comK::PactA-actA*^1-164^-*TEV-ActA*^165-639^]

ActA with TEV sequence placed between amino acids 164 and 165.

**JAT1027** [*comK: :PactA-actA*^1-196^-*TEV-ActA*^197-639^]

ActA with TEV sequence placed between amino acids 196 and 197.

**JAT1028** [*comK::PactA-actA*^1-210^-*TEV-actA*^211-639^]

ActA with TEV sequence placed between amino acids 210 and 211.

**JAT1024-1028**: The TEV protease recognition sequence (ENLYFQG) was inserted into ActA where designated. Here, the position of the TEV recognition sequence within ActA is relative to the start codon. In the manuscript, these constructs are labeled as TEV33, TEV109, TEV134, TEV166 and TEV180. These numbers correspond to the number of amino acids after the signal peptidase cleavage site and not the number of amino acids after the start codon.

**JAT1037** [*Pspa-spa*^1-51^-*nsp*^173-603^-*actA*^600-639^]

*Pspa:* protein A (*spa*) promoter. SpA^1-51^: protein A (SpA) signal sequence.

**JAT1040** [*tRNA^Arg^: :Pspac/lacOid-actA*^1-639^]

*Pspac/lacOid:* SPAC/lacOid IPTG-inducible promoter.

For strains JAT817 and JAT818, constructs were inserted into the previously described non-integrating, expression vector pHCMC03^54^. For strain JAT1037, the non-integrating vector pOS1 was used^56^. The remaining strains used integrating vectors pPL1 or pPL2^55^, as described in the Strain construction **section**.

